# A lottery simulation model for predicting cancer risk based on the strength and abundance of individual mutations

**DOI:** 10.1101/087916

**Authors:** Alexander J Wolfe, Adam Giangreco

**Affiliations:** Lungs for Living Research Centre, University College London, 5 University Street WC1E 6JF

## Abstract

Models for predicting an individual’s risk of developing cancer are complicated by a lack of well-conserved mutations present in most types of disease. To address this challenge we developed a simple lottery simulation of cancer development based solely on the abundance and relative survival advantage of mutations present within individual stem or progenitor cells. In simulations requiring few mutations or that involve mutations exhibiting a strong survival advantage cancers develop in our simulation in a progressive manner amenable to oncogene-based risk prediction. In contrast, in simulations involving numerous mutations that lack a strong survival advantage, a stochastic process of neutral drift and punctuated equilibrium determines eventual tumour formation. In these situations, the development of cancer is largely attributable to the rate of mutational variability amongst progenitor cells rather than the identity and relative survival advantage of the mutations present at any single time point. These results suggest that measuring rates of stem cell mutational variability represents an important component of predicting an individual’s risk of developing cancer.

## INTRODUCTION

The theory that cancers are the end-result of accumulated oncogenic changes was first posited by Nordling, Armitage and Doll based on their observations regarding carcinogen dose responsiveness, tumour latency, and age-dependent mortality (Nordling 1953, Armitage and Doll 1954). Given enough time, individual progenitor or stem cells are thought to accumulate sufficient numbers of mutations in a stepwise manner. These progressively accumulated mutations provide cells their capacity to proliferate indefinitely, evade immunosurveillance, and invade the local tissue, thereby resulting in cancer formation (Stratton, Campbell et al. 2009). Implicit in this model is the understanding that the risk of developing most cancers depends largely on the pro-tumorigenic properties of specific oncogenic mutations and the relative survival advantage conferred by each of these changes.

In recent years, high throughput sequencing has raised the possibility of comprehensively cataloguing all mutations and chromosomal aberrations present within cancers (Stratton, Campbell et al. 2009, Pe’er and Hacohen 2011). The aim of these studies has been to identify conserved genetic mutations suitable for predicting cancer risk and facilitating the early clinical diagnosis of disease. Surprisingly, this work has uncovered a staggering diversity and lack of conservation amongst inter-individual and intra-individual mutations that precludes the identification of oncogenes suitable for cancer risk prediction (Wood, Parsons et al. 2007, Pleasance, Cheetham et al. 2010, Pleasance, Stephens et al. 2010). Studies have also established that the number of mutations required for cancer development varies widely depending on tissue type, and that the mutational burden within physiologically normal stem and progenitor cells is often similar to that of cancers (Ding, Getz et al. 2008, Brody 2011, Martincorena, Roshan et al. 2015). Thus, the relationship between the identity and abundance of mutations within specific tissue progenitor cells and overall cancer risk appears to be more complex than originally anticipated (Alexandrov, Nik-Zainal et al. 2013).

To address this complexity, we developed a lottery-based simulation of cancer development based on the observation that a random process of stem cell turnover governs the homeostasis of most tissues (Simons and Clevers 2011). We defined a series of randomly generated numbers, or ‘events’ as representing individual stem or progenitor cell mutations. Each of these ‘events’ confers variable degrees of selective advantage, similar to the biological variability amongst individual oncogenic mutations. To complete each simulation (equivalent to winning a lottery jackpot; here representing the development of cancer), all events must be selected. We designed our simulation to vary both the number of events required for cancer development as well as their individual survival advantage. Our simulated results provide new insights regarding the relative importance of oncogene abundance, strength, and mutation rate in human tumorigenesis.

## RESULTS

In standard lottery simulations, each winning number (event) must be chosen simultaneously and the correct sequence of these events changes after each cycle of play. In this simulation, the correct or ‘winning’ sequence of events remains a fixed constant. Additional modifications include the ability to vary the odds of a single event being chosen (representing the mutational frequency of any given gene), the ability to vary the total number of events required (representing the number of oncogenes required for cancer development), and the introduction of a ‘dwell time’, or number of plays for which each correctly chosen event persists. Dividing this dwell time over the odds of an event being chosen establishes the relative survival advantage of individual events. By varying both the total number of events and their relative survival advantage, four possible scenarios for cancer development emerge (Figure 1A).

**Figure 1.**
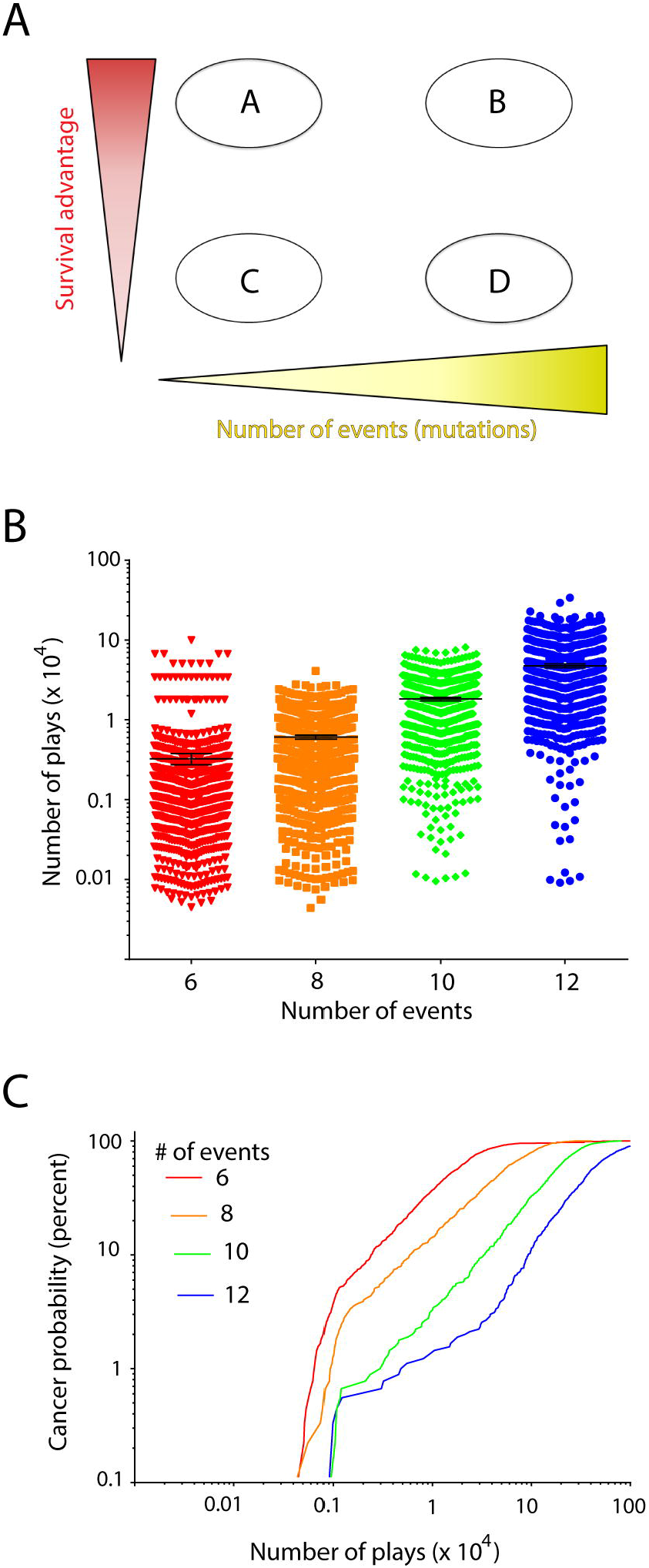
A modified lottery simulation for cancer development. (A) Varying event abundance and survival advantage produces four possible scenarios for cancer development. (B) Results of simulations requiring 6, 8, 10, or 12 total events, each of which exhibits 100% survival advantage. (C) Probability curves of cancer development versus the number of total plays for simulations requiring 6, 8, 10, or 12 events exhibiting 100% survival advantage. Error bars (B) represent the standard error of the mean; each simulation was run 1000 times to minimize sampling variability.

To test our model, we examined whether changes to the number of events influenced the number of ‘plays’ necessary to complete each simulation. This was done under conditions in which correctly selected events were maintained for a dwell time equal to the odds of each event occurring (100% survival advantage; scenarios A and B, Figure 1A). Using these parameters, we observed a positive correlation between the number of events and the number of plays required (Figure 1B). Each simulation was run 1000 times, and we plotted the number of plays required to complete each simulation against the proportion of total simulations (Figure 1C). These results resembled an approximation of the age-dependent incidence of the 4 most common adult human cancers (breast, prostate, lung, colorectal) as reported by the SEER database (http://seer.cancer.gov). Thus, our simulation adequately reproduces the natural kinetics of human tumour formation.

We next studied whether varying the survival advantage of individual events influenced the number of plays required to complete each simulation. We examined simulations requiring either 6 or 12 total events and varied the survival advantage of individual events between 10% -100% (6 events, scenarios A versus C, Figure 1A) or 40%-100% (12 events, scenarios B versus D, Figure 1A). We also introduced a ‘powerball’ event that, once present, persisted for the entirety of each simulation. This was done to mimic the occurrence of fundamental driver mutations (such as p53) that are present within the majority of human cancers and thought to influence subsequent oncogene accumulation within affected cells. Consistent with our previous results, we observed an inverse correlation between the relative survival advantage of individual events and the number of plays required to complete each simulation (Figure 2A, B). Interestingly, although varying the relative survival advantage of individual events increased the number of required plays, it had no appreciable effect on the kinetics of each simulation (Figure 2C, D). Thus, we were unable to distinguish which of these simulated scenarios might best reflect natural human tumorigenesis. There was also no increase whatsoever in the rate of cancer development following the introduction of a ‘powerball’ event, consistent with the observation that p53 and other driver mutations are frequently observed at high abundance within otherwise physiologically normal tissues (Martincorena, Roshan et al. 2015).

**Figure 2.**
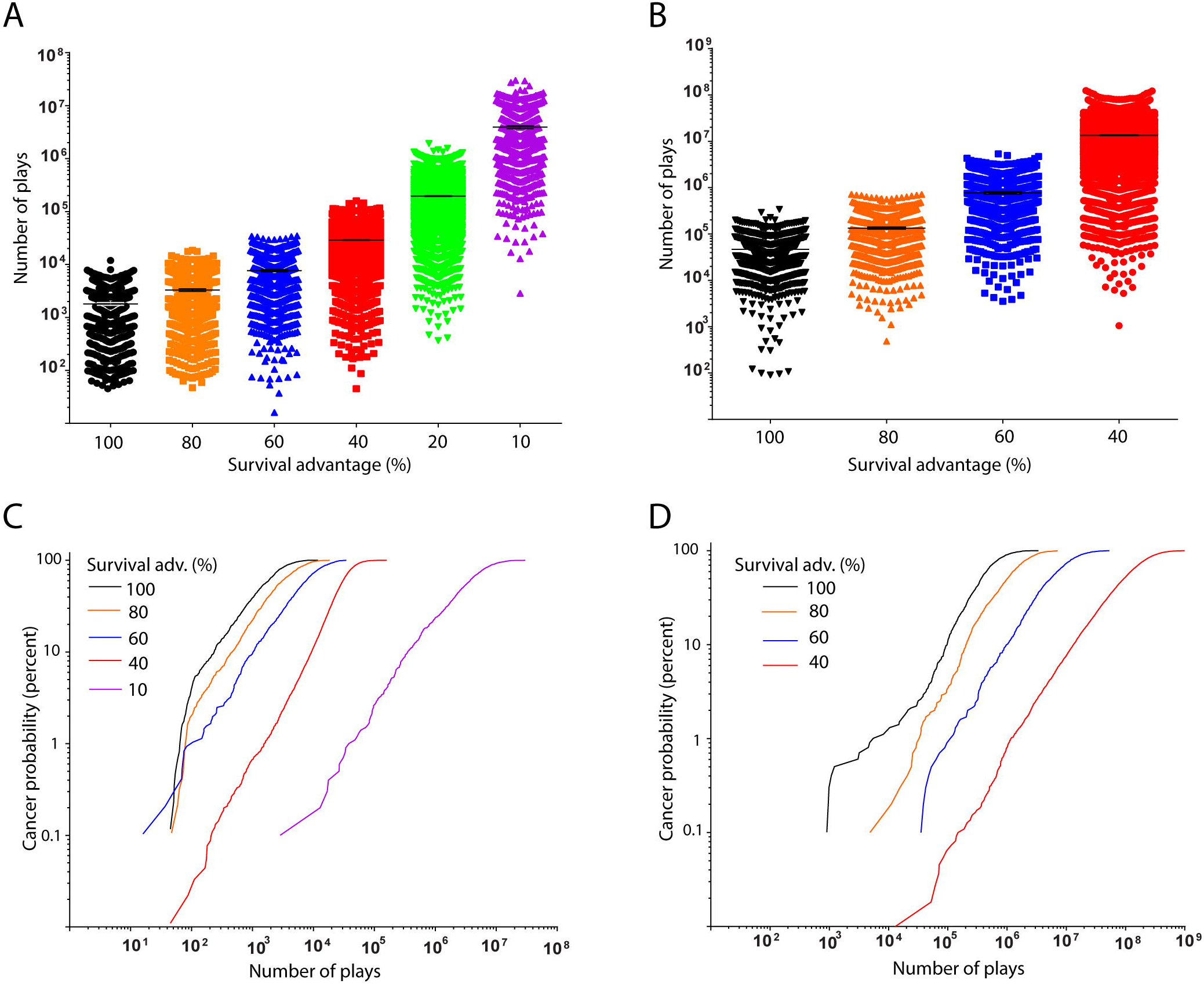
Reducing the survival advantage of individual oncogenes slows the rate of cancer development. (A, B) Results of simulations requiring 6 (A) or 12 (B) total events, each of which exhibit between 10-100% survival advantage as indicated. (C, D) Probability curves of cancer development plotted against the number of plays requiring 6 (C) or 12 (D) total events with varying survival advantage. Error bars (A, B) represent the standard error of the mean; each simulation was run 1000 times.

In simulations requiring either 6 or 12 events, we examined the rate and pattern of individual event accumulation. In simulations where events exhibited a 100% survival advantage, we invariably observed a linear, direct relationship between event accumulation and the number of plays required to complete each simulation (Figure 3A, B). Similarly, we found that under conditions of moderate (40-60%) survival advantage scenarios requiring 6 total events exhibited a progressive, linear event accumulation (Figure 3C). In contrast, under conditions of moderate survivial advantage the accumulation of events in scenarios requiring 12 total events occurred in a random, stochastic manner (Figure 3D). A similar pattern in simulations involving 6 total events was observed under conditions of low (<20%) survival advantage (data not shown).

**Figure 3.**
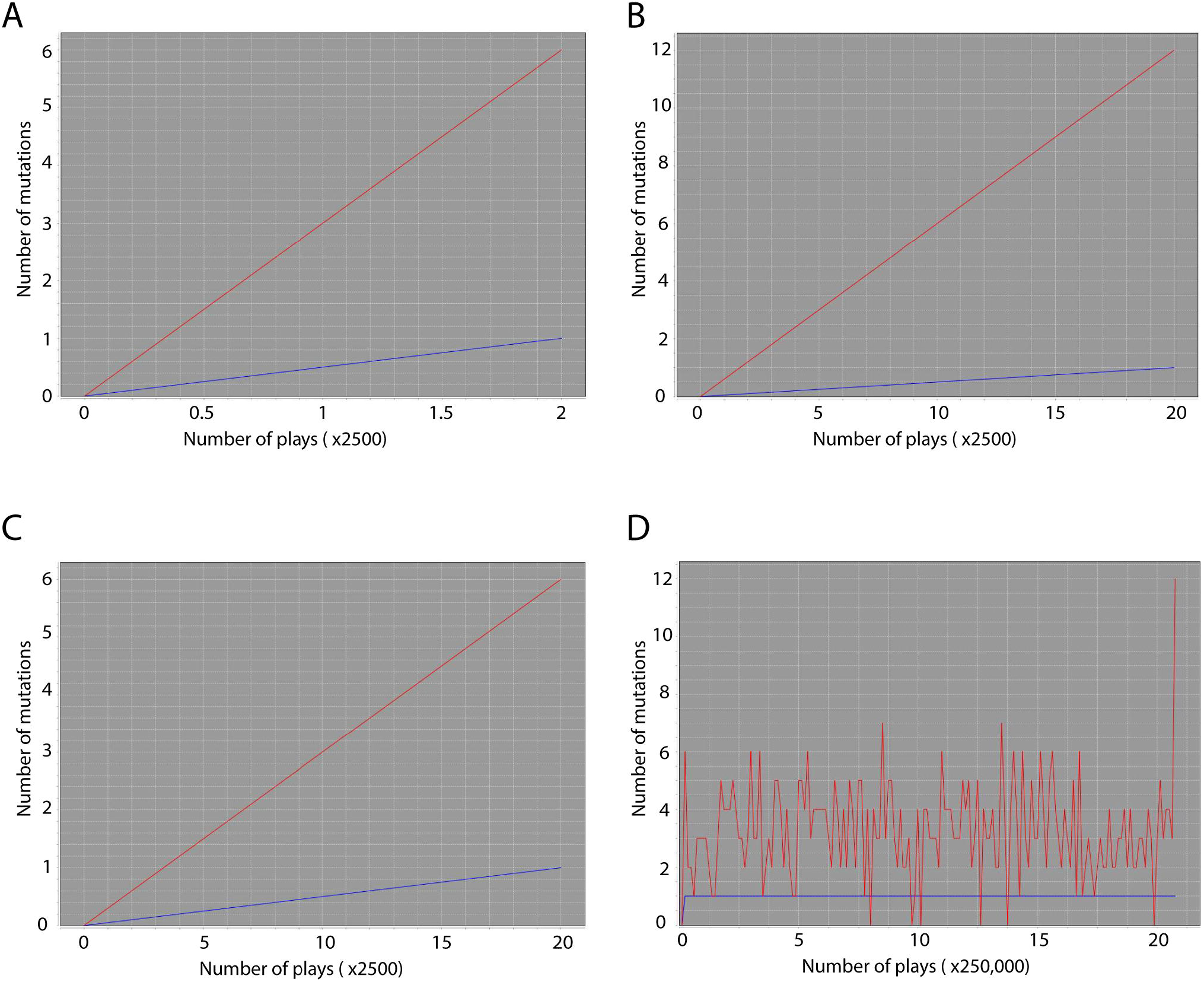
The pattern of oncogene accumulation varies based on the abundance and survival advantage of individual mutations required for cancer development. (A, B) Plots of individual simulations requiring 6 (A) or 12 (B) events, each of which exhibits 100% survival advantage. (C, D) Plots of simulations requiring 6 (C) or 12 (D) total events, each of which exhibits 40% survival advantage. Total event abundance is represented by a red line; ‘powerball’ events by a blue line (A-D).

## DISCUSSION

The present study uses a simple lottery simulation to provide new theoretical insights regarding the role of oncogene strength, abundance, and mutation rates in determining the probability of developing cancer. An important component of this simulation is that it does not require *a priori* knowledge of the identity of specific oncogenes associated with individual cancers. Rather, the only information needed is an approximation of the survival advantage conferred by individual progenitor cell mutations and the number of mutations required for initiating tumour formation. In simulations that require few mutations or that involve mutations exhibiting a strong survival advantage (scenarios A-C, Figure 1A) we found that cancers develop in a progressive manner amenable to oncogene-based risk prediction. In contrast, we observed that in simulations involving numerous oncogenic mutations that lack a strong survival advantage (scenario D, Figure 1A), a stochastic process of mutational variability is likely to precede tumour formation. Using these data, we establish that in addition to the strength of individual oncogenic mutations, the rate of mutational variability within individual progenitor cells is an important, previously unrecognised predictor of cancer risk. Our data also suggest that, while undoubtedly important, the presence of single driver mutations such as p53 is unlikely to influence the kinetics of tumour development. Furthermore, our results support the recently described ‘big bang’ scenario of tumorigenesis, in which a process of punctuated equilibrium is thought to facilitate the accumulation of significant mutational burdens within cells over a short space of time.

In order to determine the clinical utility of our simulation it will be necessary to establish the identity of mutations present within longitudinally collected precancerous biopsy samples. The goal of such a study will be to identify whether oncogenic mutations accumulate in a progressive, stepwise fashion amenable to gene-based risk prediction (scenarios A-C, Figure 4) or whether mutations occur in a random, unpredictable fashion with a high degree of temporal variability (scenario D, Figure 4). Scenarios A through C, in which oncogenic mutations accumulate linearly with time, would facilitate oncogene based risk stratification and early patient diagnosis. This is likely to be the case for cancers with highly conserved oncogene profiles such as childhood leukemias. In contrast, should scenario ‘D’ be observed, this would suggest that these tumours are driven by a stochastic process of punctuated equilibrium resulting in the progressive accumulation and subsequent disappearance of numerous ontogenic mutations. In this case, an assessment of an individual’s cancer risk based on individual oncogenes is unlikely to be effective. Rather, predicting cancer risk in these patients may only be achieved via an analysis of the rate of mutational variability. This possibility is consistent with the recent observation that multiple oncogenic mutations are present within hundreds of mutated clones throughout physiologically normal skin and oesophagus tissue (Martincorena, Roshan et al. 2015). This scenario is also in agreement with the surprisingly high number of mutations and lack of oncogenic conservation present within most types of cancer (Wood, Parsons et al. 2007, Pleasance, Stephens et al. 2010, Pe’er and Hacohen 2011).

**Figure 4.**
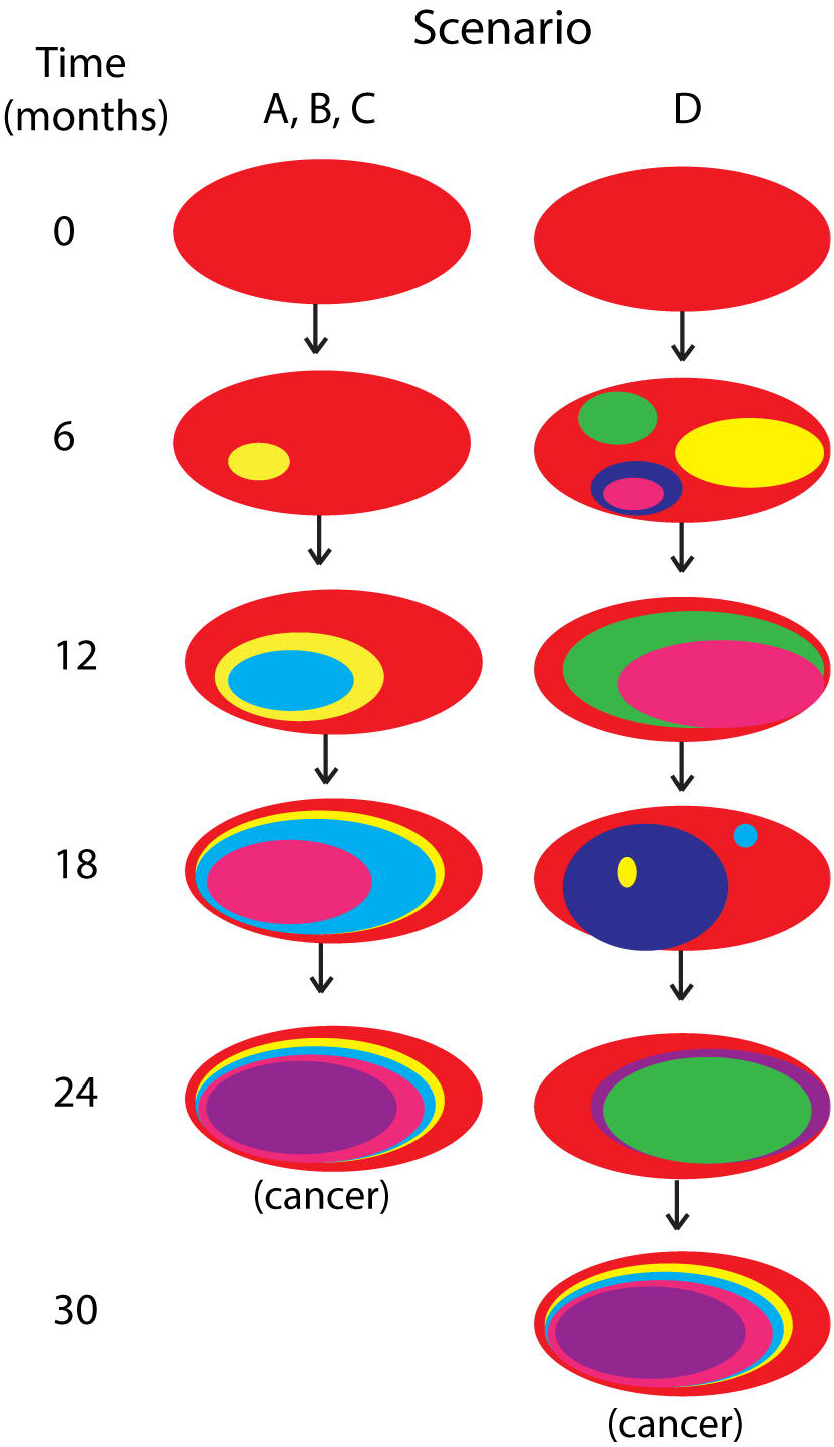
Depictions of theoretical progressive versus stochastic mutation accumulation in precancerous biopsy specimens. Scenarios A-C depict the accumulation of mutations in a progressive, stepwise fashion amenable to oncogene-based risk prediction. Scenario D depicts the random, unpredictable fashion with which mutations are expected to occur in cancers requiring numerous oncogenes that lack a significant survival advantage.

Overall, our simulation predicts that, given adequate time, all proliferative tissues exhibiting imperfect DNA replication will eventually develop cancers. Our model also predicts that the mechanisms by which cancers arise depend primarily on the strength and number of oncogenic mutations. These findings extend the work of Tomasetti and Vogelstein regarding stem cell division and cancer risk and support their suggestion that the most promising avenue for reducing cancer risk is through prevention (Tomasetti and Vogelstein 2015). Specifically, our results indicate that strategies specifically aimed at reducing rates of mutational accumulation and variability over time will reduce overall cancer incidence. These strategies will inevitably involve reducing cellular turnover, toxicant and mutation exposure via dietary and lifestyle modification. Thus, contrary to the rules of a normal lottery game, in our simulation ‘the only way to win (i.e., prevent cancer development) is not to play’.

## Acknowledgments

We gratefully acknowledge Professors Sam Janes, Benjamin Simons, and Peter Campbell for assistance in the conceptualization and interpretation of these results. We acknowledge Jeff Miller and Jason Seba for help with designing the stochastic lottery simulation model, and members of the Lungs for Living Research Centre for helpful comments regarding data interpretation. Dr Adam Giangreco is supported by a European Research Council Starting Investigator Grant (#260290) and the UCL Comprehensive Biomedical Research Centre.

